# Effects of *Bacillus thuringiensis* subsp. *kurstaki* application on non-target nocturnal macromoth biodiversity in the eastern boreal forest, Canada

**DOI:** 10.1101/2024.07.19.604320

**Authors:** Jodi O. Young, Joseph J. Bowden, Eric R. D. Moise, Robert Scott, B. Christian Schmidt

## Abstract

Insect biodiversity is crucial for resilient ecosystems, supporting essential services. Moths play significant roles as herbivores, pollinators, decomposers, and as prey for birds, bats, and predatory invertebrates, emphasizing their conservation importance. However, large-scale studies on the non-target effects of *Bacillus thuringiensis* subs. *kurstaki* (Btk) on moth communities are limited, particularly in northern forests where diversity is lower. Btk is commonly used to control pest insect populations, with forest managers in eastern Canada applying it to manage eastern spruce budworm.

We established a replicated two-year study in the North American Boreal Forest of western Newfoundland, Canada, using paired Btk-treated and control (untreated) sites.

No significant differences in total abundance or richness were found between treated and control sites. In 2021, Hill numbers only differed between our northern treatment and control sites, which may reflect those stands having already received multiple years of treatment compared to just one year in the southern sites. In 2022, control sites showed higher diversity (Shannon and Simpson diversity metrics) compared to treated sites, extending to all locations.

Multiple years of Btk treatment led to shifts in community composition and the relative abundance of some common species, without affecting total richness or abundance. Species responses varied, likely due to Btk sensitivity, application timing, and differences in phenology and voltinism, making it difficult to generalize the effects of Btk on moth communities.

## Introduction

Insect biodiversity is vital to all ecosystem services such as supporting services (e.g., decomposers), provisioning services (e.g., pollinators), regulation services (e.g., serving as a food source), and cultural services (e.g., ecotourism) (Sanmartín-Villar & Cordero-Rivera, 2024). The order Lepidoptera (moths and butterflies) is highly diverse and well represented in boreal forests. Investigating the biodiversity patterns and dynamics of Lepidoptera species offers crucial insights into the complexity of ecosystem health and resilience. While butterflies are more well known, moths are 10 times more taxonomically diverse (Wagner *et al*., 2021). They provide numerous important ecosystem services such as defoliation, pollination, nutrient cycling, plant decomposition, and prey for other animals (Holmes, Schultz & Nothnagle, 1979; Hammond & Miller, 1998; Wagner, 2005). Nocturnal moths have been shown to be more efficient pollinators of certain plants, such as bramble (*Rubus fruticosus* agg.) (Anderson, Rotheray & Mathews, 2023). Moths also differ substantially in their flight phenology, meaning that early season flyers are likely to pollinate a different cohort of plants than those that emerge later. Moths are an important food source for many animals such as predatory invertebrates, birds and bats (Wagner *et al*., 2021) Still, relatively little is known about the status of this largely nocturnal insect group, and to conserve the species and the important ecosystem services they provide we must have data on the current state of moth biodiversity and the potential impacts of forest management techniques.

The Tortricidae is a family of micromoths that contains numerous species of economic and ecological importance in forests. One of them, the spruce budworm (*Choristoneura fumiferana*, Clem., henceforth SBW), has a particularly large impact on fir and spruce forests across Canada and the United States (Maclean & Ostaff, 1989; Chang *et al*., 2012). SBW outbreaks occur cyclically approximately every 30-40 years and often last for many years (Blais, 1983). During these outbreaks, stands are heavily defoliated which leads to reduced growth and eventual tree mortality (MacLean, 1984). Within the 20^th^ century, there were three major outbreaks in Canada, resulting in tens of millions of hectares of damage (Blais, 1983). These outbreaks have both economic (e.g., timber losses, Chang *et al*., 2012) and ecological consequences (e.g., changes to wildlife habitat and carbon sequestering ability; Dymond *et al*., 2010; Chagnon, Bouchard and Pothier, 2022), therefore appropriate management strategies are imperative for forest health.

*Bacillus thuringiensis* subsp. *kurstaki* (*Btk*) has been widely used in Canada to manage this irruptive pest since the mid-1980s (van Frankenhuyzen, 2000). *Btk* is a naturally occurring soil bacterium that produces proteinaceous crystals, endowing it with insecticidal properties(Lambert & Peferoen, 1992). *Bacillus thuringiensis* subsp. *kurstaki* is a bacteria comprised of spores that produce a crystalline toxin specifically harmful to Lepidoptera (Schünemann, Knaak & Fiuza, 2014). Once consumed by larval stages, the toxins contained within the crystals are released in the gut of the caterpillar and bind with specific receptors in the lining, ultimately resulting in their death by cell lysis (Knowles & Ellar, 1987; Lambert & Peferoen, 1992). *Btk* is applied aerially onto forest stands intending to protect the current year’s foliage and/or reduce the outbreak spread and magnitude (Fuentealba et al., 2019; Johns et al., 2019). Previous studies have shown that *Btk* is an effective insecticide for lepidopterous pests (Bauce *et al*., 2004; Sanahuja *et al*., 2011). With *Btk* being Lepidoptera specific, it can negatively impact non-target moth species.

Previous studies have shown an overall negative effect on abundance and richness of non-target species (Boulton et al., 2002; Boulton, 2004; Miller, 1990; Sample et al., 1996).

However, other research has shown that the impact on non- target species is not linear and taxa are not equally affected, with differences in phenology and reproductive traits playing a role (Boulton et al., 2002; Boulton & Otvos, 2004; Leza et al., 2021; Manderino et al., 2014; Miller, 1990, 2000; Peacock et al., 1998; Strazanac et al., 2005; Wagner et al., 1996).

Moreover, there is evidence to suggest that *Btk* may have a positive effect on some genera due to decreased competition in sprayed areas (Sample *et al*., 1996; Manderino, Crist & Haynes, 2014). The variation in susceptibility, in combination with the rarity of large-scale field experiments, makes it difficult to predict the non-target effects of *Btk* application *in vivo* and highlights the knowledge gaps that still exist.

The objective of this study was to quantify *Btk* effects on non-target macromoth species in the context of a SBW management program. Forest management practices that negatively impact lepidopteran communities can have significant trophic implications across all levels of the food chain, by either decreasing prey availability to higher trophic levels or modifying the richness of consumers of primary production (Summerville, 2011), all of which pose ecosystem consequences. As one of the most functionally significant taxa in forest ecosystems, nocturnal macromoths were selected for this study due to their diversity, abundance, and ease of capture and identification (Holmes, Schultz & Nothnagle, 1979; Burford, Lacki & Covell, 1999; Young, 2005; Alison *et al*., 2022).

Using a replicated ‘treatment vs. control’ design, we made use of an ongoing “Early Intervention Strategy” (Johns et al., 2019) established to manage SBW in western Newfoundland, Canada, to investigate how the aerial application of *Btk* in forests affects the community structure (i.e., diversity, abundance, and composition) of nocturnal macromoths.

## Materials and Methods

### Study Location

We conducted this study over two years (2021 and 2022) on the west coast of Newfoundland, Canada. The study sites were divided into two locations: north and south of Gros Morne National Park, Newfoundland and Labrador, Canada (Figure 1). The northern sites were located in the Northern Peninsula Forest ecoregion (Protected Areas Association of Newfoundland and Labrador, 2008a). These sites experience cool summers and prolonged winters, characterized by some of the coldest temperatures on the island. The annual rainfall in these regions ranges between 1300-1500mm. Climate modeling using the CanESM2 model revealed average temperatures for June, July, and August to be 11.2°C, 16.8°C, and 17°C, respectively (Chylek *et al*., 2011). The forested areas in which we sampled are dominated by balsam fir (*Abies balsamea*, (L.) Mill).

**Figure 1.**
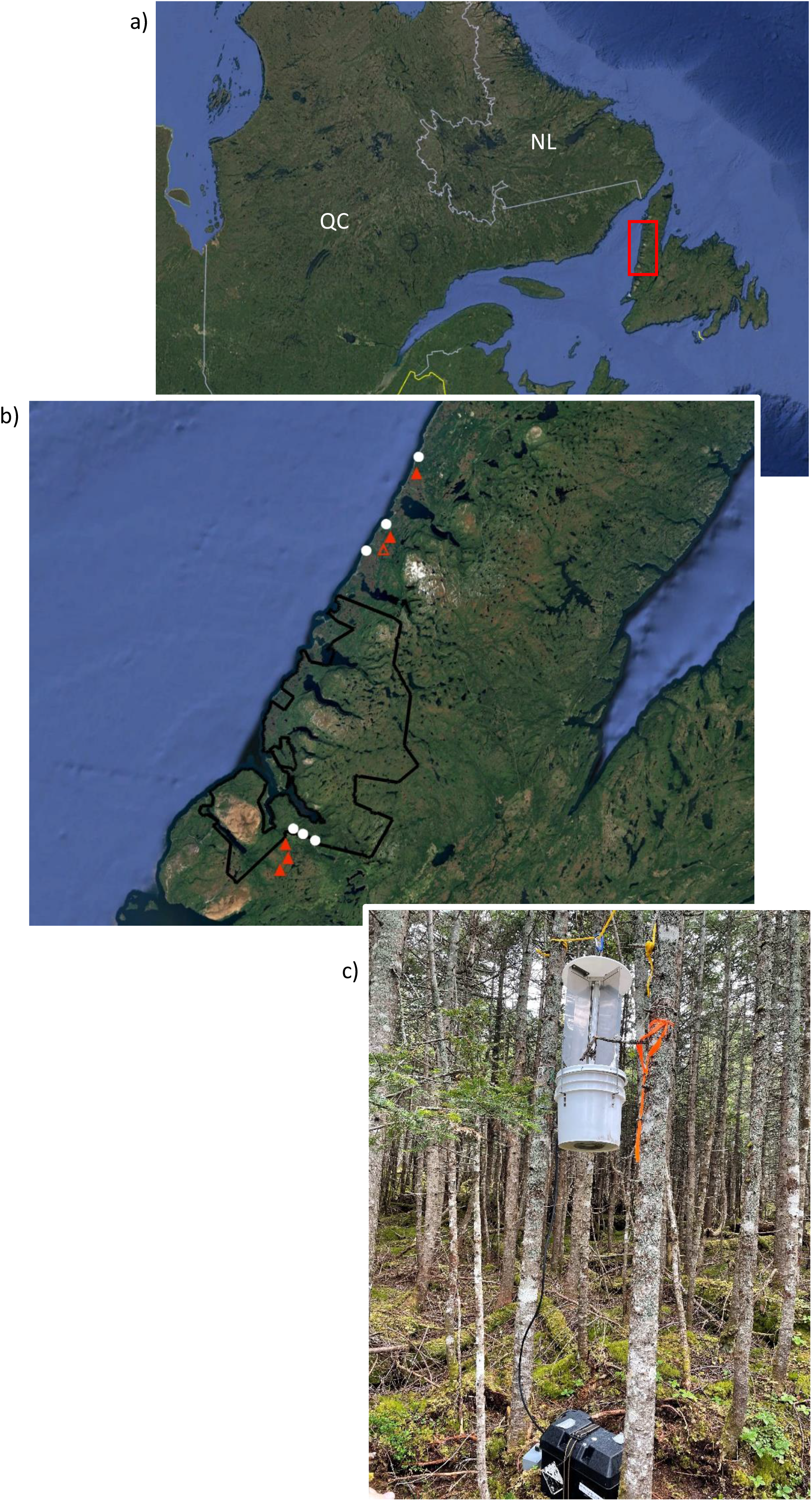
a) Newfoundland, Canada. b) Region of study in western Newfoundland. White circles represent control (non-Btk treated) sites, orange triangles represent Btk treated sites, and the hollow orange triangle represents the replaced site. c) Black-light fluorescent trap set up for moth collection at one of the boreal forest sites.

The southern sites were located in the Western Newfoundland Forest ecoregion (Protected Areas Association of Newfoundland and Labrador, 2008b). These sites experience warm summers and cold winters with an annual rainfall of 1200mm. Based on CanESM2 climate modelling, the average temperatures for June, July, and August are 12°C, 17.1°C, and 16.9°C, respectively. Consistent with the northern sites, all collections took place in balsam fir dominated boreal forest stands.

We sampled a total of 13 sites. In 2021, we collected moths from 12 sites in a paired design of treatment and control sites. In the south, there were three sites that had been sprayed with *Btk* for one year (2021) and three control sites that had not been sprayed. In the north, there were three sites that had been sprayed for two consecutive years (2020 and 2021) and three control sites that were not sprayed. In 2022, we sampled most of these sites again, with an extra consecutive year of *Btk* treatment: in the south, the treatment sites had been sprayed for two consecutive years (2021 and 2022) and in the north, the treatment sites had been sprayed for three consecutive years (2020, 2021, and 2022). The control sites remained the same. In 2022, one northern treatment site had to be shifted to a nearby stand and replaced as it was no longer included in the 2022 spray program, and we wanted to maintain consistent spray parameters across all sites. We chose a new site approximately 4.5 kilometers away from the previous site. As with all northern treatment sites, this one had been sprayed for three consecutive years (2020, 2021, and 2022).

### Spray Program

The Spruce Budworm Early Intervention Strategy is a management strategy adopted by the Healthy Forest Partnership, a group of federal, provincial, academic, and industry stakeholders. The goal of this program is to identify potential SBW outbreak “hotspot” locations by employing landscape-level monitoring, then proactively curtailing the growth of SBW populations with application of *Btk* before insects surpassing outbreak thresholds. The province of Newfoundland and Labrador began their treatment in 2020, actioned by their Department of Fisheries, Forestry, and Agriculture (Government of Newfoundland and Labrador, 2020).

*Btk* was used in 2020, 2021, and 2022 to protect 35 583, 138 950, and 142 474 hectares of forested land, respectively (Government of Newfoundland and Labrador, 2020; https://www.gov.nl.ca/ffa/programs-and-funding/forestry-programs-and-funding/idc/controlprogram/). This largely took place in the Great Northern Peninsula using one or more applications at a concentration of 30 billion international units per liter (BIU/L) at a standard rate of 1.5 liters per hectare (L/ha) for a total of 45 BIU/ha. All aerial applications were applied when weather conditions and larval development were optimal for *Btk* treatment. These optimal weather conditions were considered when: wind speeds were between 2 and 10km/h, air temperatures were below 25°C, relative humidity was above 50%, and it was not raining or expected to rain within 2 hours of treatment (Government of Newfoundland and Labrador, 2019). In 2020, spraying took place between June 26^th^ and July 16^th^. In 2021, all spraying took place between June 7^th^ and July 12^th^. Finally, in 2022, all spraying took place between June 13^th^ and July 8^th^. In 2020, all spraying was limited to areas north of Gros Morne National Park; consequently, our study sites south of the park received one year less treatment than the northern sites.

### Moth collection

At each site we used one bucket light trap to capture adult nocturnal macromoths. Each trap was equipped with a 12W black-light fluorescent tube light that was powered by a deep-cycle lead-acid battery (Figure 1C). 5-gallon buckets were attached to hold collected moths and three insecticide strips as a killing agent (Hercon® Vaportape II™). The traps were also equipped with photocell light sensors that turned the tube lighting on at dusk and off at dawn.

We placed the traps approximately two meters off the ground by tying them to nearby trees. Sampling took place over two summers (2021 and 2022), with three collections (June, July, and August) per year. Although this does not encompass the breeding times of all species, this is when the majority are active (Pinksen *et al*., 2021). We chose sampling dates based on the lunar cycle, with the new moon being targeted due to a reduced influence of moonlight (Yela & Holyoak, 1997).

Sampling in 2021 took place June 15-17, July 13-15, and August 10-12, and in 2022 from June 28-30, July 27-29, and August 23-25 for a total of 108 trap-nights per year. There was no trap disturbance during sampling. During each sampling event, the traps were set up for three days and emptied daily into collection boxes that allowed for adequate air flow to promote drying and limit mold growth. Once collected, samples were sorted to retain only moths that belonged to families that are considered macromoths (i.e., Drepanidae, Geometridae, Saturniidae, Sphingidae, Notodontidae, Erebidae, and Noctuidae). Moths were collected and identified to the lowest possible taxon using a variety of resources (Beadle & Leckie, 2012; *Moth Photographers Group*, 2022; John, 2022) A voucher collection was created and is stored with Natural Resources Canada, Canadian Forest Service, Corner Brook, Newfoundland and Labrador, Canada. This voucher collection was confirmed by B. C. S.

### Statistical analyses

We completed all data analysis using R (version 4.2.2; R Core Team, 2023). We separated the data by stand type (North Control, North Treatment, South Control, South Treatment). Separate analyses were conducted in 2021 and 2022.

To assess changes in moth abundance, we used a generalized linear model using the *glm.nb* function in the R package *MASS* (Ripley *et al*., 2023), fitted with a negative binomial distribution. The model assessed the impact of stand type on absolute moth abundance per site, pooling data across months to reduce the effect of seasonal variation. Singletons were excluded, and post hoc analyses using pairwise Tukey’s tests were conducted for significant findings between treatment types. The top five most abundant species per year, constituting ≥35% of the collection each year, were similarly analyzed. For each stand, diversity indices, encompassing species richness, Shannon diversity, and Simpson diversity, were calculated using Hill numbers (q = 0; species richness, q = 1: Shannon diversity, q = 2; Simpson diversity) with the *iNext* package (Chao *et al*., 2014; Hsieh, Ma & Chao, 2016). To visualize differences in species composition between stand types, nonmetric multidimensional scaling (NMDS) ordination was calculated with Bray-Curtis dissimilarity scores using the function *MetaMDS* in the R package *Vegan* (Oksanen *et al*., 2019). A species-by-site matrix was created using transformed counts (x’=log(x+1)). Data was pooled by month and singletons were removed to eliminate the influence of rare species. The ordination was plotted using the function *ggplot* in the R package *ggplot2* (Wickham, Navarro & Pedersen, 2016). Two dimensions were chosen as it was promoted visualization and interpretation. To assess differences between stand types, we performed a permutational multivariate analysis of variance (PERMANOVA) using the function *adonis2* in the R package *Vegan* (Oksanen *et al*., 2019).

## Results

We collected and identified 11,802 macromoths, representing 174 species that belonged to seven families (Drepanidae, Erebidae, Geometridae, Noctuidae, Notodontidae, Nolidae, and Sphingidae). Of these, 95.7% were identified to species level with the remaining unidentifiable.

### Abundance

In 2021 and 2022, overall moth abundance was significantly influenced by stand type (p<0.001 and p=0.0208, respectively; Table 1, Figure 2). In 2021, no significant differences were observed between treatment and control groups within the north or south regions, the south control sites had higher abundance than the north control or north treatment sites (p<0.001 and p=0.0121, respectively). Although not statistically significant, the mean abundance trended higher in the north treatment group than the north control group and in the south control group than the south treatment group. Like the previous year, we detected no significant difference between the treatment and control groups within the north and south regions in 2022. The only significant difference was between the north control sites and south control sites, with more individuals in the south control sites (p=0.0467).

**Figure 2.**
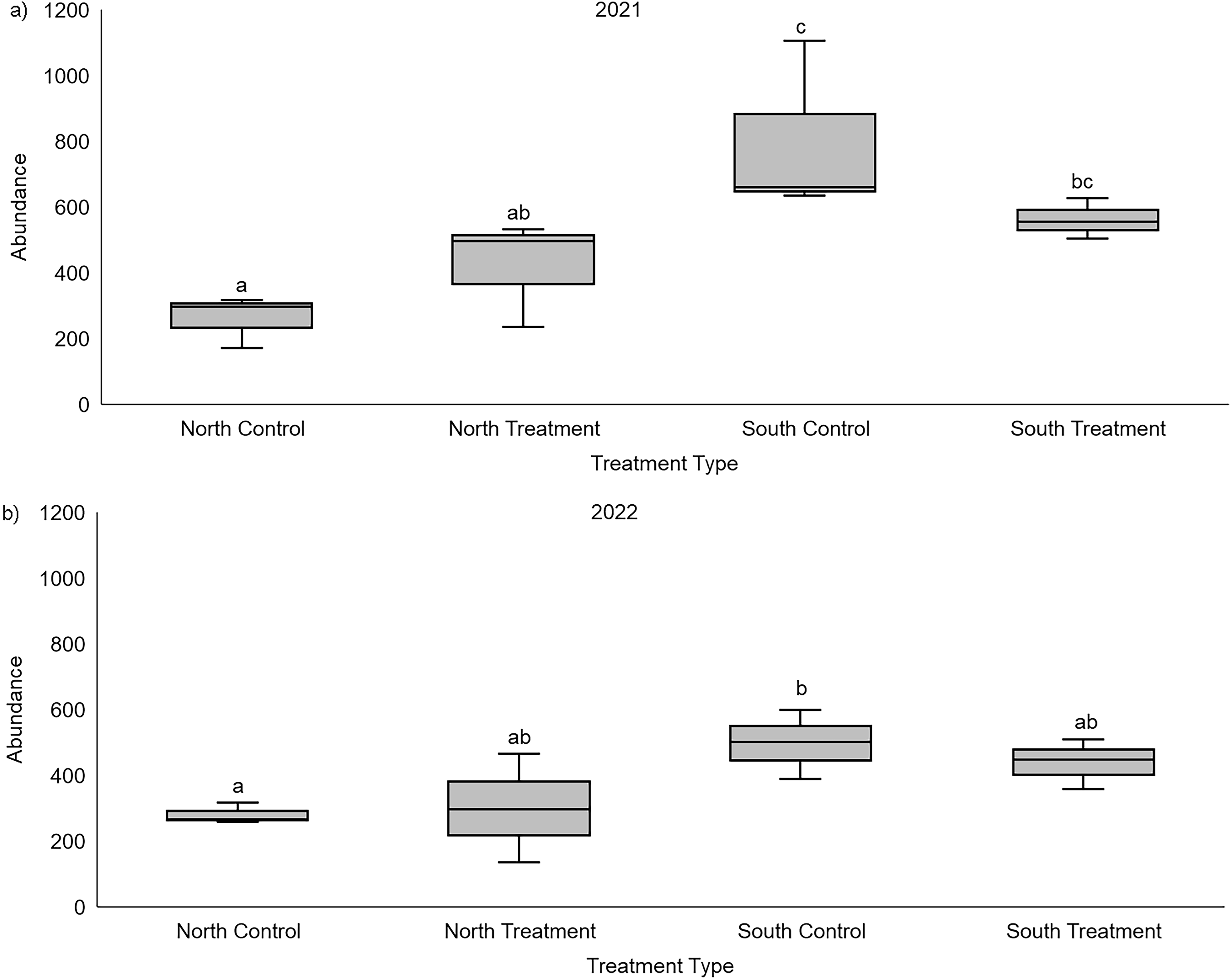
Box and whisker plots of moth abundance from four stand types, across two years, a) 2021 and b) 2022. 2021 treatment sites had been sprayed with *Btk* for one (South Treatment) and two (North Treatment) consecutive year(s). 2022 treatment sites had been sprayed with *Btk* for two (South Treatment) and three (North Treatment) consecutive years. Comparisons tested using a generalized linear model (GLM), fitted with a negative binomial distribution. Letters a, b, and c indicate significant differences between stand types based on post-hoc testing (95% confidence).

**Table 1.**
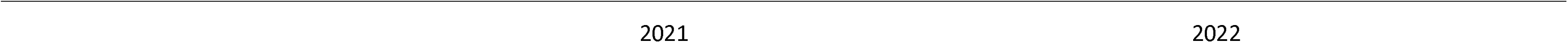
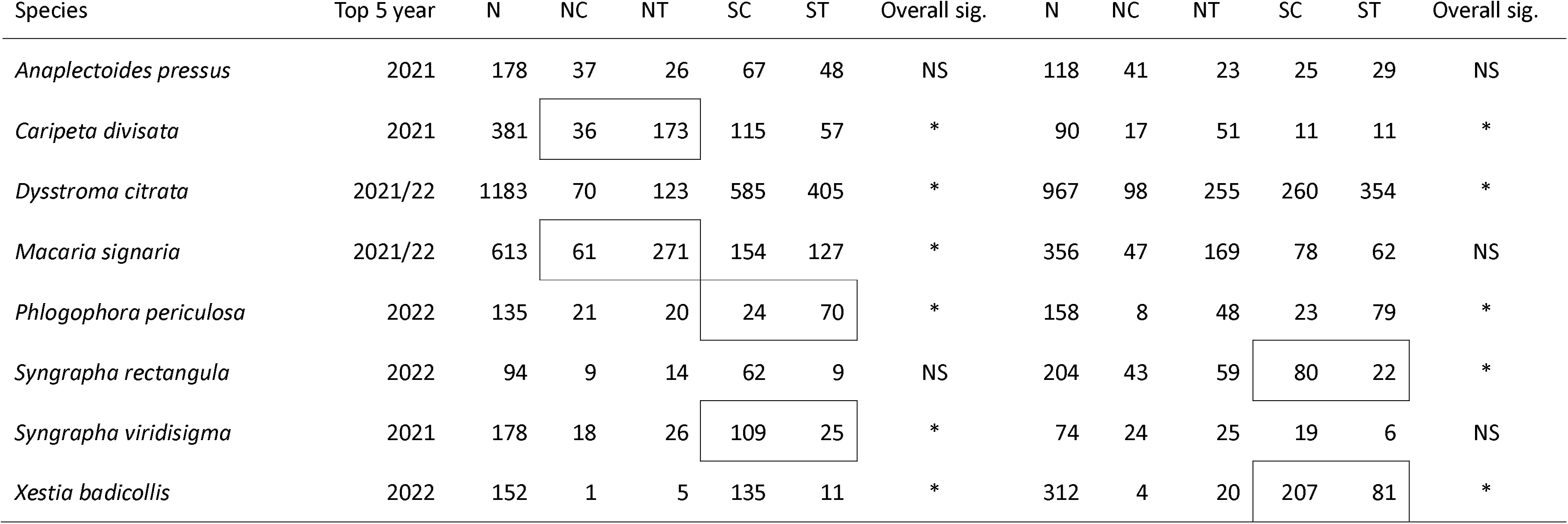
Total raw moth abundance for the five most abundant species in 2021 and 2022. Collection days amalgamated by year and stand type. Control/treatment pairs that appear in a box are significantly different. NC = North control, NT = North treatment, SC = South control, ST = South treatment

However, the mean abundance was higher in the north treatment group than in the north control group and higher in the south control group than in the south treatment group.

We assessed any changes in abundance in the most common species across both years. In 2021, the five most abundant species made up 39% of the collection, with *Dysstroma citrata* (Linnaeus; Geometridae) (n=1183), *Macaria signaria* (Hübner; Geometridae) (n=613), *Caripeta divisata* (Walker; Geometridae) (n=381), *Syngrapha viridisigma* (Grote; Noctuidae) (n=178), and *Anaplectoides pressus* (Grote; Noctuidae) (n=178). In 2022, the five most abundant species accounted for 35% of the collection for that year. Two species remained the same as 2021 (*Macaria signaria* (n=333) and *Dysstroma citrata* (n=869)), while three were different (*Xestia badicollis* (Grote; Noctuidae) (n=152), *Syngrapha rectangula* (Kirby; Noctuidae) (n=94), and *Phlogophora periculosa* (Guenée; Noctuidae) (n=133)). We detected significant differences in the abundance of these species among stand types in both 2021 and 2022.

In 2021, significant differences were detected in the abundance of *Syngrapha viridisigma, Macaria signaria, Caripeta divisata, Phlogophora periculosa, Dysstroma citrata,and Xestia badicollis* among stand types (p<0.001 each). *Syngrapha viridisigma* were more abundant in the south control sites than the south treatment sites (p=0.0273), with no significant difference observed between treatment and control in the north. *Macaria signaria* was significantly more abundant in north treatment sites than in north control sites (p = 0.0221), while no significant differences were found in the south.

*Caripeta divisata* was more abundant in north treatment than north control (p < 0.001), and *Phlogophora periculosa* was more abundant in south treatment than south control (p < 0.001). *Dysstroma citrata* and *Xestia badicollis* had an overall significant difference in abundance detected but no pairwise difference was detected. In 2022, *Xestia badicollis*, *Syngrapha rectangula, Caripeta divisata*, *Dysstroma citrata*, and *Phlogophora periculosa* showed significant differences in abundance based on stand type (p < 0.001 for all except p = 0.0213 for *S. syngrapha*), with controls having higher abundance. More *Xestia badicollis* individuals were collected in south controls than south treatments (p = 0.0487). Although not significant, there were more individuals in north treatment than north control. For *Syngrapha rectangula*, a significant difference was observed between south control and south treatment sites (p = 0.013), with higher numbers in controls. Despite not being statistically significant, north control had more individuals compared to north treatment. The other three species did not have significant pairwise results.

### Diversity

In 2021, we observed significant differences in Shannon diversity (q=1) and Simpson diversity (q=2) between the north control and treatment sites (Figure 3), but not in species richness (q=0). There were no differences between the south control and treatment sites. Despite higher species richness in the controls, confidence intervals overlapped between north control and treatment, as well as between south control and treatment, indicating no significant differences.

**Figure 3.**
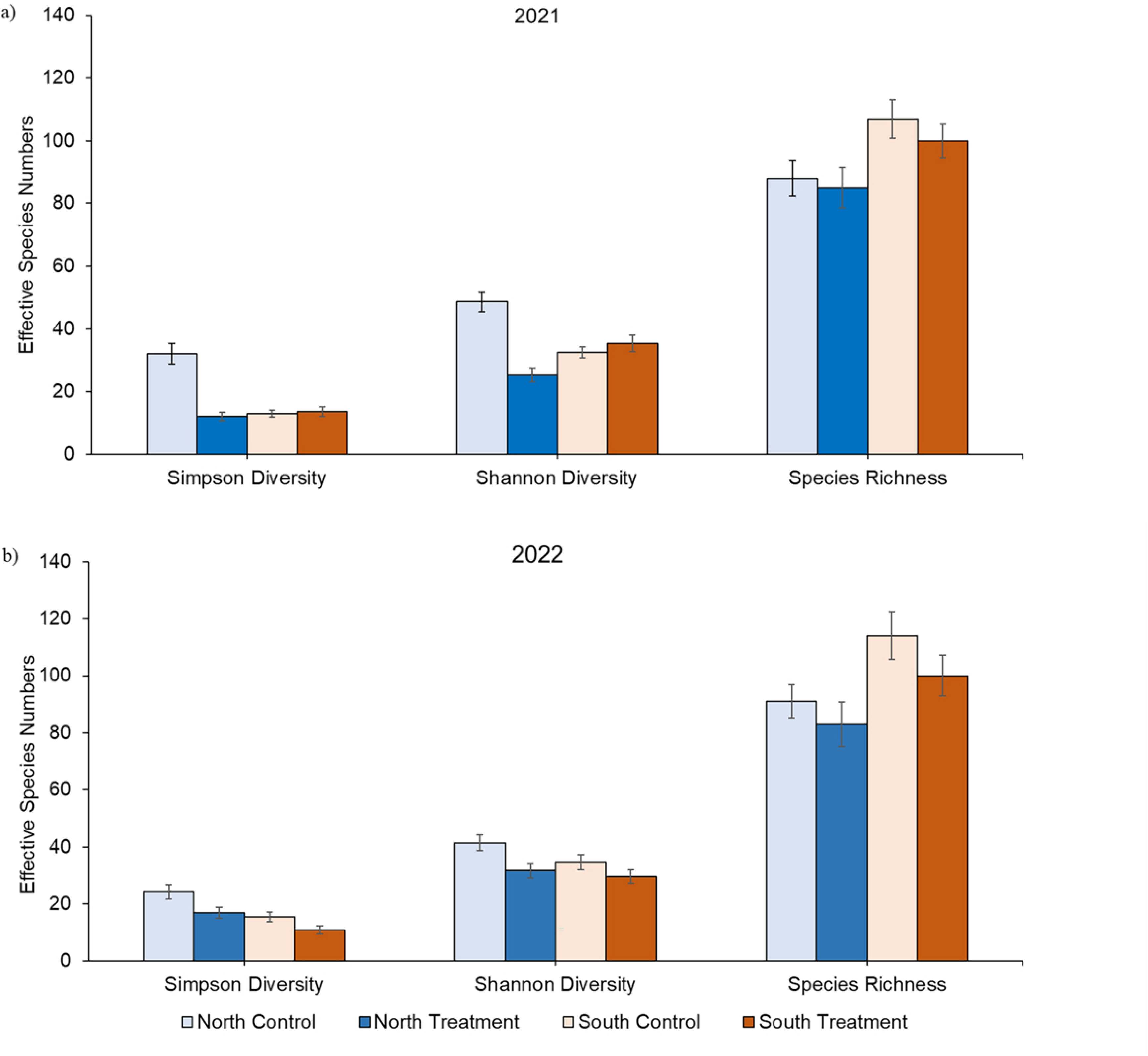
Diversity indices, characterized by effective number of species (± 95% confidence interval), for each treatment group for a) 2021 and b) 2022. Values calculated from Hill number order q = 0 (species richness), q = 1 (Shannon diversity), and q = 2 (Simpson diversity), for each stand type.

Shannon diversity was similar in the south control and south treatment sites, while confidence intervals did not overlap between the north control and north treatment sites, suggesting higher diversity in the north control. Simpson diversity was similar in the south control, south treatment, and north treatment sites, but higher in the north control sites.

In 2022, Shannon diversity (q=1) and Simpson diversity (q=2) differed between stand types, with non-overlapping confidence intervals between the north treatment and controls, and the south treatment and controls. Although species richness (q=0) had overlapping confidence intervals, it was higher in the north controls than the north treatments and higher in the south controls than the south treatments.

For Shannon diversity, there was no overlap in confidence intervals between the south control and south treatment sites, as well as between the north control and north treatment sites, indicating higher diversity in the controls in both regions. Simpson diversity was higher in the north control group than the north treatment group and higher in the south control group than the south treatment group.

Between years, we note differences in Simpson and Shannon diversity, but not in species richness based on overlapping confidence intervals. Simpson diversity increased in the north treatment sites and decreased in the control sites from 2021 to 2022, with no differences found in the southern sites. Shannon diversity increased in the north treatment sites but decreased in the south treatment and north control sites. No difference between years was observed in the south control sites.

### Species composition

NMDS ordination indicated that species composition varied among stand types (Figure 4). The visualization of the ordination was recapitulated statistically by the PERMANOVA results (2021: F=2.1505, p=0.002; 2022: F=2.7568, p=0.001; stress < 0.1 for all). However, pairwise post-hoc testing did not identify significant differences between any specific pairs of groups. The 2021 ordination plot revealed distinct clustering patterns, with the south treatment group located in the bottom right quadrant, the south control group located in the top right quadrant. The northern treatment and control groups overlapped in the upper and lower left quadrants; however, they were distinct from the southern groups. Similarly, the 2022 ordination revealed distinct clustering, with the south control group being in the lower right quadrant, the south treatment group being in the upper right quadrant, the north control being in the lower left quadrant, and the north treatment being in the upper left quadrant.

**Figure 4.**
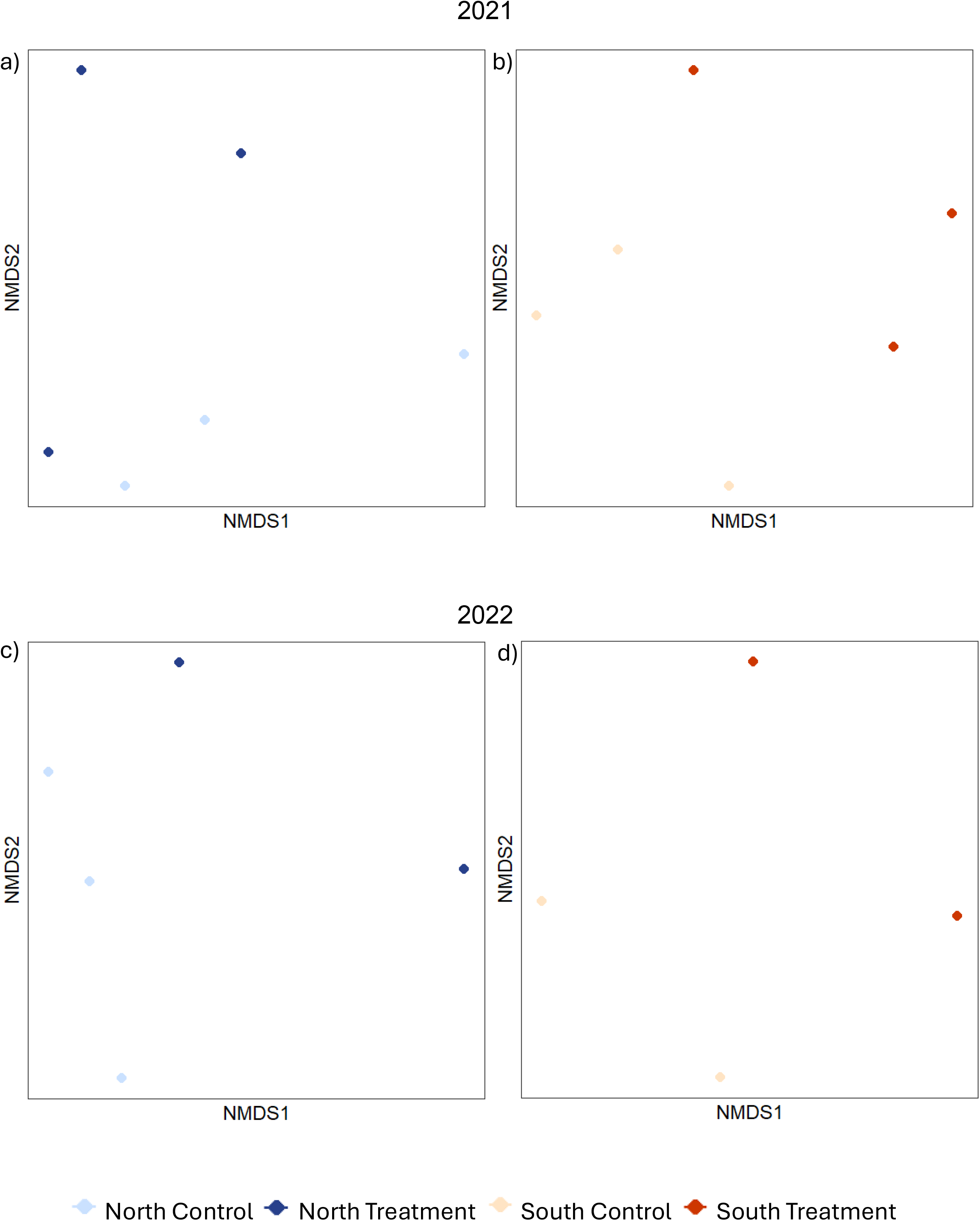
Nonmetric multidimensional scaling of species composition across stand types in a) 2021(Stress=0.119) and b) 2022 (Stress=0.093). Species abundance pooled across all dates and log transformed (x′ = log(x+1)). Bray-Curtis dissimilarity scores were used. Each individual treatment plot represents a collection month for north control, north treatment, south control, and south treatment.

### Discussion

Our results show that overall macromoth abundance and richness was not higher in the control sites, but community composition did differ between sites treated with Btk and those that were not (controls). Patterns of changing relative abundance among species were reflected in both the Shannon and Simpson diversity indices. Although species richness did not differ significantly, it was consistently higher in control sites. These complex results point to the need for a more nuanced and thorough assessment of interspecific variation in *Btk* impacts on non-target Lepidoptera, with particular emphasis on understanding individual species responses to *Btk* treatment that could reveal deeper phylogenetic patterns of susceptibility.

### Abundance

We found that the overall abundance of macromoths did not differ between *Btk* treated and control (untreated) sites. These findings are consistent with Glaus *et al*. (2023), who employed similar *Btk* application rates. However, while both of our studies found that the overall abundance of non-target Lepidoptera did not differ between *Btk-treated* areas and non- treated areas, our present study found, although not always significant, there is evidence of interspecific variation in treatment effects. For instance, in 2021 we found that six of the eight most common species across both years (*Dysstroma citrata, Macaria signaria*, *Caripeta divisata, Syngrapha viridisigma*, *Xestia badicollis* and *Syngrapha rectangula*) were more abundant in the north treatment than the north control.

This pattern then carried over to the 2022 season where seven of the eight most common species were more abundant (*Caripeta divisata*, *Dysstroma citrata, Phlogophora periculosa*, *Syngrapha rectangula Xestia badicollis)*. In the south sites, only one of the most abundant species (*Phlogophora periculosa*) was higher in the treated sites in 2021, but this increased to three species (*Anaplectoides pressus*, *Dysstroma citrata,* and *Phlogophora periculosa)* in 2022.

Our findings are in contrast to those of Boulton (2004) who observed a decline in non-target caterpillar abundance after *Btk* application. In their study, their *Btk* application was only for one year, but the spray rate was higher than in our current study. Sample et al. (1996) also found a decrease in abundance of both adult and larvae non-target macromoths following *Btk* application. However, their application was significantly less than ours: further highlighting the complex susceptibility of non-targets. Our results were influenced heavily by the five most abundant species in each year, which accounted for 39% and 35% of total macromoths in 2021 and 2022, respectively. These species were the primary drivers of total macromoth abundance. This is in contrast to the findings of Wagner *et al*. (1996) who found the abundance of the most common species to be decreased in the treated sites, although his findings were not statistically significant. Moreover, they treated their plots with one application of 90 BIU/ha while our sites were treated with 45 BIU/ha. This discrepancy between the findings of Wagner et al. (1996) and our study highlights the potential importance of the concentration (and possibly frequency) of *Btk* application. The applications of lower doses may have allowed for reduced immediate impacts and potential recovery periods, contributing to differing outcomes in our study. These distinctions underscore the importance of considering dosage frequency and intensity in understanding nuanced effects of *Btk* application on non-target moth populations. While our findings reveal no significant differences in overall abundance, this is attributed to the notable increase observed in less than 40% of our entire collection each year; therefore, the remaining 60% offsets any potential discernible variations in the total moth population.

Because species differ in phenology, their respective exposure to Btk can differ substantially. Among Lepidoptera taxa, sensitivity to *Btk* application often aligns with life history characteristics (Peacock et al., 1998). Species with an early phenology, are multivoltine, or univoltine with a long larval period appear to be less sensitive to *Btk* (Boulton, Otvos & Ring, 2002; Boulton, 2004). Although comprehensive phenological data are lacking for all prevalent species in this study, some patterns emerge. For instance, the activity window of *Macaria signaria*, which spans from May to August, suggests that it would have surpassed its larval stage prior to *Btk* application. *Anaplectoides pressus*, *Caripeta divisata*, and *Syngrapha rectangula*’s emergence around June aligns with their pupal stage occurring prior to *Btk* application. In contrast, the flight times of *Dysstroma citrata*, *Syngrapha viridisigma*, *Phlogophora periculosa,* and *Xestia badicollis* begin in July, potentially coinciding with the SBW’s flight. The overlap in flight times between these and the SBW could suggest that they have overlapping larval stages, meaning they likely would be feeding during *Btk* applications. However, we did not find the abundance of these species to be lower in the treated sites. This opens another avenue of investigation for other possible explanations of why some species may be more-or-less susceptible to *Btk*.

Certain species may inherently exhibit reduced sensitivity to *Btk* (Peacock *et al*., 1998), potentially contributing to the absence of significant differences in abundance between treatment and control sites. This may explain why *Dysstroma citrata*, *Syngrapha viridisigma*, *Phlogophora periculosa,* and *Xestia badicollis* were slightly more abundant in the treatment sites. Given the limited understanding of these dynamics, this presents an interesting avenue for future investigation.

### Diversity

In the north sites, our findings indicate different outcomes between 2021 and 2022. Both years showed greater Shannon and Simpson diversity in control groups compared to treatment groups, with consistently higher richness in controls, though not significantly different. This suggests a shift in the relative abundance of different species within the community in treated sites, leading to uneven species distribution and ultimately creating diversity differences. The analysis of species with low and intermediate abundance revealed minimal differences between treatment and control groups. However, focusing on dominant species, including *Anaplectoides pressus*, *Caripeta divisata*, *Dysstroma citrata*, *Macaria signaria*, *Phlogophora periculosa, Syngrapha rectangula, Syngrapha viridisigma*, and *Xestia badicollis* were more abundant in the north treatment than the north control, influencing evenness and explaining the diversity difference. The control sites exhibited a higher diversity due to a more equitable distribution of abundance across species.

In the south, few differences in richness and diversity between treatment and control groups were observed in 2021. However, in 2022, diversity decreased in treatment sites. In 2021, only one dominant species (*Phlogophora periculosa*) was more abundant in treatment sites, but in 2022, three of eight dominant species (*Dysstroma citrata*, *Anaplectoides pressus*, and *Phlogophora periculosa*) showed higher abundance in treatment sites. Richness remained similar among stand types in 2021 but was slightly higher in control sites. In 2022, Shannon and Simpson diversity were greater in the control group than the treatment group, while richness was similar. This suggests that shifts in relative abundance, especially among dominant species in treatment sites, account for variation in diversity.

2021 was the first year of treatment in the southern sites, followed by a second application in 2022. Lack of diversity difference in 2021, followed by a decrease in 2022, suggests a lagged effect of adult macromoth responses to *Btk* application. This aligns with previous studies noting significant impacts occurring 12 to 14 months post-spray (Sample *et al*., 1996; Boulton *et al*., 2007).

### Species composition

We anticipated differences in species composition between treated and control sites due to documented changes in relative abundance following exposure to *Btk* (Wagner et al., 1996). Our ordination analysis revealed distinct clustering patterns, indicating differences in species composition between treated and untreated stands.

While PERMANOVA analysis revealed significant differences in species composition among stand types, there were no significant pairwise differences. This suggests that while overall differences exist in moth community structure in response to *Btk* treatment, these differences may not manifest uniformly at the level of individual pairwise comparisons between treated and untreated (control) sites.

The lack of significant pairwise differences underscore the complexity of community responses to *Btk* treatment and suggests that the impacts may vary across different species or sites. Despite the non-significance of the pairwise comparisons, the significant overall differences indicated by PERMANOVA suggest that *Btk* treatment exerts discernible effects on the composition of moth communities, with potential implications for ecosystem dynamics.

## Conclusions

It is a rare and unique opportunity to study the impacts of insect management at regional spatial scales in forests. Here, we have shown that the application of *Btk* to control the current SBW outbreak in western Newfoundland significantly affected moth species community structure in the first few years of SBW management. While the overall differences in moth community composition were significant, the impacts of treatment may not be consistent across all regions. Moreover, macromoth responses to *Btk* are somewhat idiosyncratic in that interspecific variation in developmental phenology and treatment sensitivity is likely to influence overall impacts; however, understanding such mechanisms will require further research.

Although significant, impacts observed in our study were much less than those reported for treatments of longer durations and higher application rates (Wagner *et al*., 1996; Boulton, 2004). Minimizing such effects will require a balance between SBW outbreak control while minimizing *Btk* application, which has been successfully achieved elsewhere (MacLean *et al*., 2019). The current treatment program was initiated in the context of the Spruce Budworm Early Intervention Strategy framework (Johns *et al*., 2019), and non- target impacts will ultimately depend on successful execution of this proactive program. With Lepidoptera providing important ecosystem services such as herbivory, pollination, and a food source for other species (Hammond & Miller, 1998), it is imperative to understand the non-target impacts of forest pest control programs. Such information will allow landscape managers to develop strategies that effectively balance pest control measures with the preservation of biodiversity in boreal forest ecosystems.

## Acknowledgements

We would like to thank Memorial University – Grenfell Campus for financial support. We would also like to thank Parks Canada, Gros Morne National Park for access to some sites for data collection (permit: GMP-2022-42098 to JJB). We would also like to acknowledge the collaborative efforts of the Dept. of Fisheries, Forestry and Agriculture, Newfoundland and Labrador as we would not be able to conduct this large-scale field experiment without their cooperation. Lastly, we would like to thank Jamie Warren for his technical help.

